# Priming of C-glycoside flavones in *Colobanthus quitensis* with salicylic acid, methyl jasmonate, pimelic acid, suberic acid, and azelaic acid elicits antifungal activity against *Botrytis cinerea*

**DOI:** 10.1101/2025.03.30.646165

**Authors:** Lydia Rubilar, Javiera Avilés, Michelle Sarmiento, Felipe Sobarzo, Gustavo E. Zúñiga, Rodrigo A. Contreras

**Affiliations:** Laboratorio de Fisiología y Biotecnología Vegetal, Facultad de Química y Biología, Universidad de Santiago de Chile. 3363, L. B. O’Higgins Ave., Estación Central, Santiago, Chile

**Keywords:** *Colobanthus quitensis*, elicitors, C-glycosyl flavones, priming, natural antifungals

## Abstract

*Colobanthus quitensis*, one of only two native angiosperms in Antarctica, produces C-glycosyl flavones with antifungal activity against *Botrytis cinerea*. In this study, the exogenous application of the elicitors salicylic acid (SA), methyl jasmonate (MeJA), pimelic acid (PA), suberic acid (SuA), and azelaic acid (AzA) was evaluated for their effect on the accumulation of bioactive metabolites in *in vitro*-cultivated plants. Exposure to these compounds significantly modulated the expression of key genes in the phenylpropanoid and flavonoid pathways, including *pal*, *chs*, *chi*, *fnsII*, as well as regulatory genes such as *myb12*, *bhlh*, and *wrky33*, enhancing PAL activity and the accumulation of schaftoside, neoschaftoside, saponarin, and swertiajaponin. This priming process improved the antifungal activity of the extracts, with MeJA and PA identified as the most effective treatments. The *in vitro* culture approach enabled the assessment of a protected and hard-to-access species without the need for wild harvesting. These results suggest that the exogenous application of elicitors constitutes an efficient strategy to modulate the biosynthesis of specialised metabolites, with implications for the development of biocontrol agents and the improvement of efficiency in sustainable agricultural systems.

**GRAPHICAL ABSTRACT:** 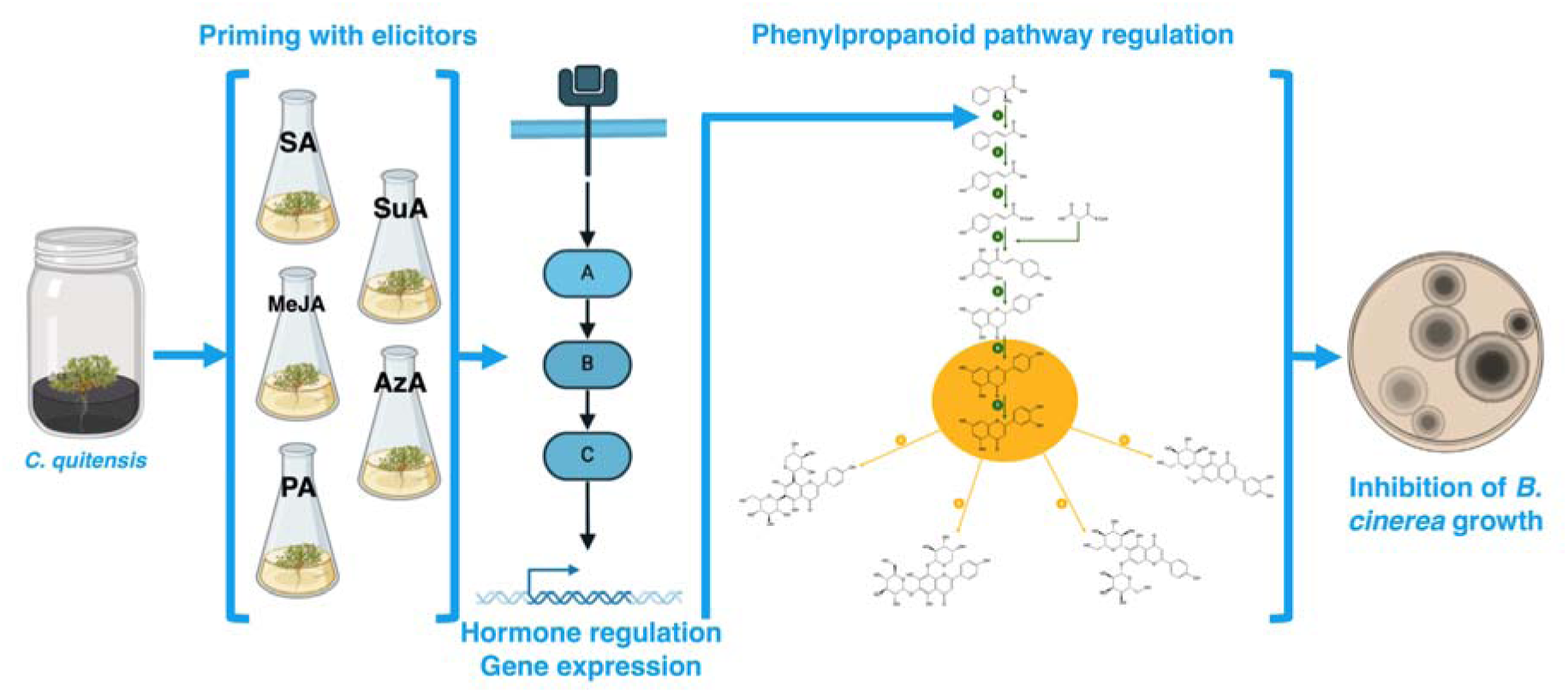

## 1. INTRODUCTION

*Colobanthus quitensis* (Kunth) Bartl. is one of only two native angiosperms in the maritime Antarctic region, making it an attractive model for studying plant adaptations to extreme environmental conditions such as low temperatures, high UV radiation, and nutrient deficiency ^1^. Its ability to survive in this hostile environment has been partially attributed to the production of secondary metabolites that play key roles in defense against adverse conditions ^2^. Among these compounds, flavonoids—particularly C-glycosyl flavones— have demonstrated antifungal activity against *Botrytis cinerea*, a pathogen of agricultural significance responsible for substantial losses in horticultural crops ^3^.

The application of exogenous elicitors is an effective strategy for modulating the synthesis of bioactive metabolites in plants ^4^. Among these, salicylic acid (SA) and methyl jasmonate (MeJA) are key signaling molecules that regulate plant defense responses ^5^. SA is involved in systemic acquired resistance (SAR) and regulates the hormone accumulation and expression of genes associated with phenylpropanoid and flavonoid biosynthesis by activating phenylalanine ammonia-lyase (PAL) ^6^. Conversely, MeJA is a mediator in wound response and induced systemic resistance (ISR), activating genes linked to the production of secondary metabolites with antimicrobial and antioxidant properties ^4^.

In addition to these phytohormones, dicarboxylic acids such as pimelic acid (PA), suberic acid (SuA), and azelaic acid (AzA) have emerged as modulators of plant defense signaling ^7,8^. AzA has been shown to function as a mobile molecule that propagates systemic defense signaling in conjunction with SA, promoting the expression of pathogenesis-related (PR) genes ^8^. SuA and PA, on the other hand, may influence oxidative stress responses in plant cells and the accumulation of phenolic metabolites, indirectly modulating the activation of defense-related metabolic pathways ^9^.

Previous studies have demonstrated that the methanolic fraction of *C. quitensis* extracts inhibits both the germination of *B. cinerea* conidia and its radial growth ^3^. Furthermore, flavones such as schaftoside, swertiajaponin, and saponarin have been shown to affect the activity of fungal virulence enzymes, interfering with key processes in fungal pathogenicity ^10^. These findings suggest that the production of antifungal metabolites in *C. quitensis* is strongly regulated by hormonal signaling networks and that the exogenous application of various elicitors could enhance their accumulation.

The use of elicitors to induce the biosynthesis of natural antifungal compounds presents a yield-oriented sustainable approach to crop protection, particularly considering the increasing emergence of fungal strains resistant to synthetic fungicides and the environmental concerns associated with their use ^11^. In this context, exploring the combined effects of SA, MeJA, PA, SuA, and AzA in *C. quitensis* could provide valuable insights into the interaction of these signals in modulating the phenylpropanoid pathway and the production of flavonoids with antifungal activity.

Therefore, in this study, we propose that the exogenous application of these elicitors will enhance the accumulation of C-glycosyl flavones in *C. quitensis*, resulting in enhanced antifungal activity of its extracts against *B. cinerea*. Additionally, this approach is expected to provide new perspectives on the hormonal and metabolic regulation of plant defense in species adapted to extreme environments, with potential biotechnological applications in the production of compounds of agro-industrial interest.

## 2. MATERIALS AND METHODS

### 2.1. Plant material

*In vitro* cultivated *C. quitensis* plantlets were used in the experiment. The original plants were obtained from collections carried out on King George Island, Maritime Antarctica, under permits granted by the Chilean Antarctic Institute (INACH) in the summer of 2010 campaign at the Polish Antarctic Station, Henryk Arctowski (62°09′37″S, 58°28′24″W). For subculturing, a Murashige-Skoog (MS) medium with 3% sucrose and a pH of 4.7 ± 0.1 was used, solidified with 2.0 g × L⁻¹ of high-acyl gellan gum (Phytagel®). The medium was supplemented with 0.25 mg × L⁻¹ benzylaminopurine (BAP) and 0.1 mg × L⁻¹ naphthalene acetic acid (NAA), maintaining a 2:1 molar ratio. Each culture flask contained 15 mL of medium. Incubation conditions were set at 14 ± 2 °C, with a 16 h light / 8 h dark photoperiod ^12^. The plantlets were maintained under these conditions for 30 days, ensuring homogeneous biomass (∼5 g of living tissue per culture flask) before conducting the experiments.

After this period, the plantlets were subcultured in flasks containing the same culture medium but without gelling agents or growth regulators. They were then acclimatized for 10 days before elicitor application. The treatments included SA, MeJA, PA, and AzA at final concentrations of 10, 25, 50, 75, 100, 200, and 500 µM. The elicitors were dissolved in sterile dimethyl sulfoxide (DMSO), and the solutions were filtered using sterile PVDF filters (0.22 µm) under a laminar flow cabinet. A negative control (without elicitor) was also included. The seedlings were incubated under the same conditions as previously described for 14 days.

### 2.2. Growth and Biomass

Fresh weight (FW) was determined using an analytical balance with a precision of 0.1 mg. Relative growth (RW) was calculated using Equation (1).

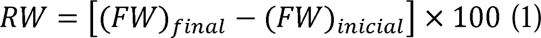

Dry weight (DW) was measured from three plantlet samples at the end of the treatment, which were dried in an oven at 70 ± 2 °C. Water content (WC) was calculated using Equation (2).

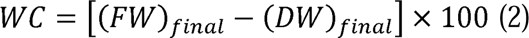

### 2.3. Metabolite extraction and analysis

#### 2.3.1. C-glycosides flavones

The extraction and quantification of phenolic compounds were performed in three independent measurements with three technical replicates each. For methanolic extraction, a proportion of 100 mg of fresh tissue was homogenized with liquid nitrogen to a fine powder and extracted in 1 mL of pure HPLC-grade methanol. The suspension was sonicated (60 Hz, 2 h, on ice), centrifuged (10,000 × *g*, 10 min, 4 °C), and the supernatant was recovered. Lipophilic fractions were consistently removed using four successive washes with n-hexane until a yellow-colored extract was obtained. Specific flavonoids, previously described in *C. quitensis*, were quantified by LC–MS/MS using an Agilent HPLC system coupled to a triple quadrupole mass spectrometer (Agilent 6410) in negative ESI mode under multiple reaction monitoring (MRM). The chromatographic separation was performed on a Zorbax Eclipse C18 column (150 × 4.6 mm, 5.0 μm) at 25 °C, using a binary solvent system composed of (A) 10 mmol × L⁻¹ ammonium acetate, pH 4.0, and (B) 10 mmol × L⁻¹ ammonium acetate in 95% acetonitrile, pH 4.0, with a flow rate of 0.6 mL × min⁻¹. The gradient elution was as follows: 0–5 min, 0–50% B; 5–10 min, 50–52% B; 10–20 min, 52–100% B; 20–35 min, 100–65% B; 35–40 min, 65–0% B ^3^. MRM transitions were used to quantify schaftoside (m/z 563.115 → 413.3), neoschaftoside (m/z 563.115 → 413.3, quantified as schaftoside equivalents), saponarin (m/z 593.166 → 431.4), and swertiajaponin (m/z 461.116 → 311.07). Data were expressed as μg × g ¹ of dry weight.

#### 2.3.2. Phytohormones

For endogenous phytohormone profiling, fresh leaves (50 mg) of *C. quitensis* were ground to a fine powder in liquid nitrogen. The homogenized samples were supplemented with a mixture of deuterated internal standards at a final concentration of 50 nM and extracted using 500 µL of a cold solution of 2-propanol, water, and HCl (2:1:0.002 v/v/v). The samples were mixed on a rotary shaker at 100 rpm for 30 minutes at 4 °C. Subsequently, 1 mL of dichloromethane was added, and the samples were centrifuged at 13,000 × *g* for 5 minutes at 4 °C. The lower organic phase was recovered and evaporated under a nitrogen stream, and the resulting residue was redissolved in 100 µL of methanol.

Quantification of phytohormones was carried out using an Agilent 6410 triple quadrupole mass spectrometer coupled to an Agilent 1260 HPLC system. Chromatographic separation was performed on a reverse-phase Zorbax Eclipse C18 column (150 × 4.6 mm, 5.0 μm), and analyte detection was achieved in multiple reaction monitoring (MRM) mode using positive or negative electrospray ionization, depending on the compound class. Specific MRM transitions were monitored for each hormone to ensure unambiguous identification and accurate quantification.

Determination was achieved by monitoring specific mass transitions for each phytohormone. The transitions included *m/z* 263.0→153.0 for abscisic acid (ABA), 209.1→59.0 for jasmonic acid (JA), 175.0→130.0 for indole-3-acetic acid (IAA), 331.1→213.0 for gibberellin A (GA), 137.0→93.0 for salicylic acid (SA), 102.0→56.0 for 1-aminocyclopropane-1-carboxylic acid (ACC), and 220.2→136.2 for trans-zeatin (tZ). All compounds were detected in negative ion mode except for tZ and ACC, which were analyzed in positive mode. Hormone concentrations were expressed as nmol × g^−1^of dry tissue ^13^.

### 2.4. Phenylalanine Ammonia-Lyase (PAL) Activity

The extraction and quantification of PAL activity were performed using three independent biological replicates, each with three technical replicates. Protein extraction was conducted by homogenizing 100 mg of fresh tissue in 1 mL of 100 mM sodium borate buffer (pH 8.8) containing 1% (w/v) polyvinylpolypyrrolidone (PVPP) and 1 mM dithiothreitol (DTT). The homogenate was centrifuged at 12,000 × *g* for 15 min at 4 °C, and the supernatant was collected for enzymatic activity assays.

Total protein quantification was performed using the Bradford method, with bovine serum albumin (BSA) as the standard. The absorbance was measured at 595 nm, and protein concentration was expressed as mg protein per gram of FW.

PAL activity was determined by monitoring the conversion of L-phenylalanine to trans-cinnamic acid following the increase in absorbance at 290 nm. The reaction mixture contained 500 μL of 100 mM sodium borate buffer (pH 8.8), 200 μL of 10 mM L-phenylalanine, and 300 μL of enzyme extract, incubated at 37 °C for 30 min. The reaction was stopped by adding 200 μL of 6 N HCl, and absorbance was recorded. A negative control was included using D-phenylalanine instead of L-phenylalanine, to confirm the specificity of the reaction.

Specific PAL activity was expressed as µkats per kilogram of protein (µkats × kg ¹ protein), where one µkat corresponds to 1 × 10 mol of trans-cinnamic acid formed per second per kilogram of protein, using a molar extinction coefficient of 10,000 M ¹ cm ¹ at 290 nm ^12^.

### 2.5. RNA extraction, cDNA synthesis, and qPCR

The total RNA extraction was performed using the Spectrum™ Plant Total RNA Kit (Sigma-Aldrich) following the manufacturer-suggested protocol. Fresh leaf tissue (100 mg) of *C. quitensis* fresh tissue was ground in liquid nitrogen and homogenized in the provided lysis solution containing 2-mercaptoethanol. The RNA was purified using the kit binding column, washed twice with wash solutions, and eluted in RNase-free water. The concentration and purity of the extracted RNA were assessed using a NanoQuant (Tecan, Austria), and RNA integrity was verified by agarose gel electrophoresis. Genomic DNA contamination was removed using the On-Column DNase I Digest Set (Sigma-Aldrich) during the extraction process.

For cDNA synthesis, 1 µg of total RNA was reverse-transcribed (RT) using the AffinityScript QPCR cDNA Synthesis Kit (Agilent) following the manufacturer guidelines. The reactions (20 µL final volume) included 10 µL of 2× First-Strand Master Mix, 3 µL of oligo(dT) primer (0.1 µg × µL^−1^), 1 µL of AffinityScript RT/RNase Block enzyme mixture, and the RNA template. The reactions were incubated at 25 °C for 5 min for primer annealing, followed by 42 °C for 15 min for reverse transcription. The enzyme was inactivated at 95 °C for 5 min, and the cDNA was stored at −20 °C until use.

Quantitative PCR (qPCR) was performed using an Agilent Stratagene Mx3005P qPCR System with Brilliant II SYBR® Green qPCR Master Mix (Agilent). The reactions were conducted in a total volume of 20 µL, containing 10 µL of 2× SYBR Green Master Mix, 0.3 µM of each primer (forward and reverse), and 1 µL of 1:10 diluted cDNA. The thermal cycling conditions included an initial polymerase activation at 95°C for 10 min, followed by 40 cycles of denaturation at 95°C for 15 s, annealing at 60°C for 30 s, and extension at 72°C for 30 s. A melting curve analysis was performed from 55°C to 95°C to ensure the specificity of amplification.

The housekeeping gene rRNA18S (*rrna18S*) was used as an internal control for normalization. Each reaction was performed in three biological replicates and two technical replicates. The relative expression levels of the target genes were calculated using the 2^(–ΔΔCt)^ method ^14^.

The following genes were analyzed, with their forward (F) and reverse (R) primers: *pal1* (F: 5’- ACACAAGAGCAACGGAGGAG -3’, R: 5’- TCCTTCTGAAGTGCGACACC -3’); *pal2* (F: 5’-ACACAAGAGCAACGGAGGAG-3’, R: 5’-TCCTTCTGAAGTGCGACACC-3’); *pal3* (F: 5’-TCCTCTCCGAGGCACCATTA-3’, R: 5’- ACCTCAATCTGCGGTCCAAG-3’); *pal4* (F: 5’-CCATCCCGGTCAGATCGAAG-3’, R: 5’-GACATGGTTGGTCACGGGAT-3’); *c4h* (F: 5’- GGAGCCGATTAGCACAAAGC-3’, R: 5’-CAATGGCGCATTTCATCCGA-3’); *4cl* (F: 5’- GTTATTTGCGTCGACTCCGC -3’, R: 5’-CGACATCGTTCGGCTGGATA-3’); *chs* (F: 5’- GCTACTCCGCCCAATGCTAT-3’, R: 5’-GACTTAGGTTGGCCCCACTC-3’); *chi* (F: 5’-AGGAGGAGGAGGAGGAGGAG- 3’, R: 5’-GGAGGAGGAGGAGGAGGAGG-3’); *fns ii* (F: 5’- CCATCACTCGCCCAAGAGTT -3’, R: 5’-TGGTATTGGCGGGTGAAGTC -3’); *ugt78d2* (F: 5’-TCCGGAGAGAAATCGCGAAG-3’, R: 5’-TCTCGATCTGAGGTTATTCGTC-3’); *ugt75c1* (F: 5’- GTCAGGTGTACCTGTGGTGG-3’, R: 5’- TTTTGCTAACTTGTAGAACAAAAGC-3’); *ugt73c6* (F: 5’- AGTTGCATCGTCATGGCTTT-3’, R: 5’-GGCAAACCAGACTCAATGGC-3’); *myb11* (F: 5’- GGGAGTTCTGATCTGCAGCC-3’, R: 5’-GATCCTGCCATTTGGCCACT-3’); *myb12* (F: 5’-TGGCCACAATGGGACGATAC-3’, R: 5’- CTGACTTGACGATATTTCTAACGAA-3’); *myb111* (F: 5’- AGTGGGGAGAGAGCAAACCT-3’, R: 5’-ACTGTTTTGTTTGGACTCAGGA-3’); *bhlh* (F: 5’-TCACCACCGTTCAATCGTGT-3’, R: 5’-CACGTGTCCACATGCAGTTA-3’); *wd40* (F: 5’-TGACTTCCTCCGCCTCTACA-3’, R: 5’-CGATACGAGACGGAACGAGG- 3’); *ttg1* (F: 5’-CCGACTCCATGTCCGTCAAA-3’, R: 5’-GGAGGGGGAGTCGGTGAT-3’); *wrky33* (F: 5’-CAAGTGTACAAGCCCCGCAT-3’, R: 5’- GTTGTGGTGGCGGTATTGG-3’); *tga2* (F: 5’-AAGCCTTGGGAGAAGGTTCC-3’, R: 5’- CTGAACTCAGGTCACGCAGG-3’); *nac42* (F: 5’- TTGGTGCCACATCTACTCATCAGG-3’, R: 5’-TGCTATTTGAACCTCCTACAACCT- 3’); *lox2* (F: 5’-CATTATCGGGCGGCCCTAAG-3’, R: 5’-CCTGTTGACATCTGGGGGAC-3’); *aos* (F: 5’-CCGACGGTGGGGAATAAACA-3’, R: 5’-GAACATGTAGAGCAGCAACAGAT-3’); *jar1* (F: 5’- TGCCACCCCAGAGATTTGCT-3’, R: 5’-TTGGCTTCCCTTGCGTAGTG-3’); *erf1* (F: 5’- CAGCGTGGTCTGAGGTGTATT-3’, R: 5’-TTGGGGGAGGAGTGAAGACT-3’); *hsr203j* (F: 5’-TTCGTGCGGTCTTATCGGAG-3’, R: 5’-ACGCTTGTTGATGAACTCTGC-3’); *pr4* (F: 5’-CCAATACGGTTGGACTGCCT-3’, R: 5’-GTGACCAGCCTGAACACCAG-3’); *rrna18S* (F: 5’-CTTAGCAGAACGACCAGCGA-3’, R: 5’- TCTTCATCGATGCGAGAGCC-3’).

### 2.6. Antifungal activity assay *in vitro*

The antifungal activity of defatted *C. quitensis* methanolic extracts standardized at 10 mg × mL⁻¹ 1 in DMSO was evaluated through radial growth inhibition assays using *B. cinerea* B05.10 strain, maintained on potato dextrose agar (PDA) plates at 22 ± 2 °C in darkness with 80% of air humidity, with final concentration of 75 µg × mL⁻¹ in each plate, based in previous published EC_50_ ^3^. The assays were performed in 6-well plates, with three biological replicates per extract and two technical replicates per sample. PDA medium was supplemented with each extract, and 5 mm diameter mycelial discs of *B. cinerea* were placed at the center of each well. The plates were incubated at 22 ± 2 °C with 80% air humidity for 72 hours in darkness, after which radial growth (mm) was measured and compared to the control to assess antifungal activity ^3^.

### 2.7. Statistical analysis

All measurements were analyzed using a two-way analysis of variance (ANOVA) followed by Tukey’s post hoc test to determine significant differences among treatments (*p* < 0.05). The ANOVA tests were performed using GraphPad Prism 10.4.1, ensuring accurate statistical comparisons. Additionally, heatmaps were generated in a Python-based environment to visualize the variation in gene expression and hormone accumulation across different treatments. The heatmaps were constructed using the Seaborn and Matplotlib libraries. Data preprocessing, including normalization and scaling, was conducted using Pandas and NumPy, while statistical significance levels were incorporated directly into the visualization for unambiguous interpretation.

## 3. RESULTS

### 3.1. Biomass production and PAL-specific activity primed by elicitors

The application of different elicitors significantly affected biomass growth and PAL activity in *C. quitensis*. In terms of biomass, growth was highest at intermediate elicitor concentrations, while higher concentrations led to a reduction. SuA induced the higher biomass increase, reaching values above 40% compared to the control at concentrations between 50 and 100 µM. MeJA and PA also promoted growth, with maximum values between 30 and 40% at 25-75 µM. In contrast, SA and AzA showed a less pronounced effect, with moderate increases at lower concentrations but a sharp decline at 200-500 µM, where biomass growth was significantly reduced compared to other treatments (Fig. 1a).

**Figure 1.**
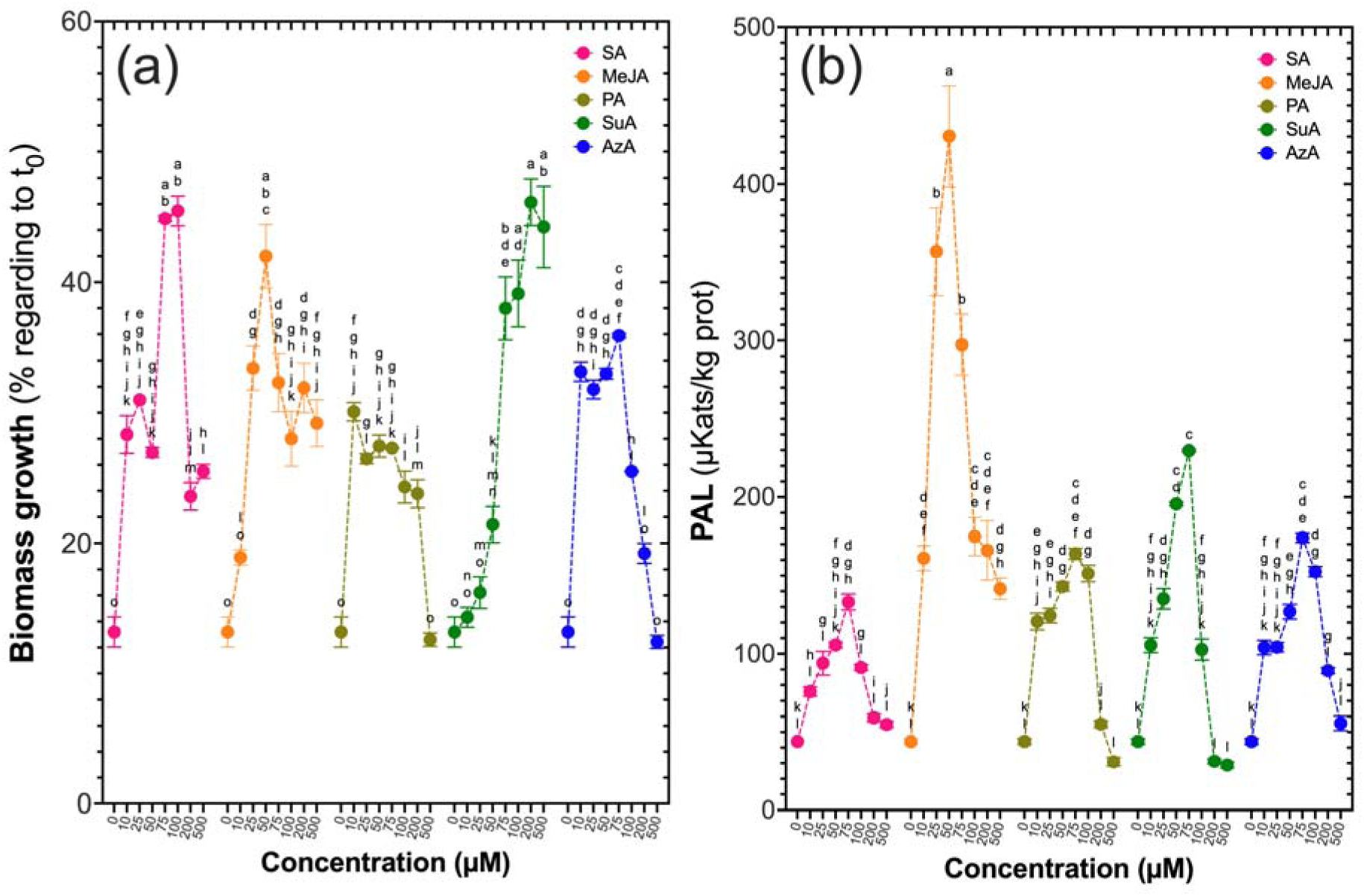
Effect of different elicitors on biomass growth and PAL activity in *C. quitensis*. (a) Biomass growth is expressed as a percentage relative to the control. (b) PAL activity expressed in µkats × kg^−1^ protein. Treatments correspond to SA, MeJA, PA, SuA, and AzA. Data represent the mean ± standard error (SE) of three biological replicates. Different letters indicate statistically significant differences among treatments (*p* < 0.05, two-way ANOVA with Tukey’s post hoc test).

PAL activity also exhibited a concentration-dependent response to the elicitors. MeJA induced the highest enzymatic activity, reaching approximately 450 µkats × kg^−1^ protein at 50 µM, followed by a gradual decrease at higher concentrations. SuA and PA also stimulated PAL activity, with peaks of approximately 300-350 µkats × kg^−1^ protein in the 50-100 µM range. In contrast, SA showed intermediate levels of induction at lower concentrations but declined beyond 100 µM. AzA exhibited the lowest PAL activation, with lower values than the other treatments and a more pronounced reduction at higher concentrations (Fig. 1b).

Overall, the results suggest that concentrations of 50-100 µM of SuA, MeJA, and PA are optimal for inducing both biomass growth and PAL activity in *C. quitensis*.

### 3.2. Phytohormone accumulation in response to elicitors

Phytohormone profiles varied according to both the type of elicitor and the concentration applied. In MeJA treatments, ABA reached its highest value at 100 µM with 27.6 nmol × g^−1^ DW, while ACC also showed a marked increase at the same concentration, reaching 33.4 nmol × g^−1^ DW. JA increased under MeJA as well, with a value of 20.9 nmol × g^−1^ DW at 100 µM (Fig. 2).

**Figure 2.**
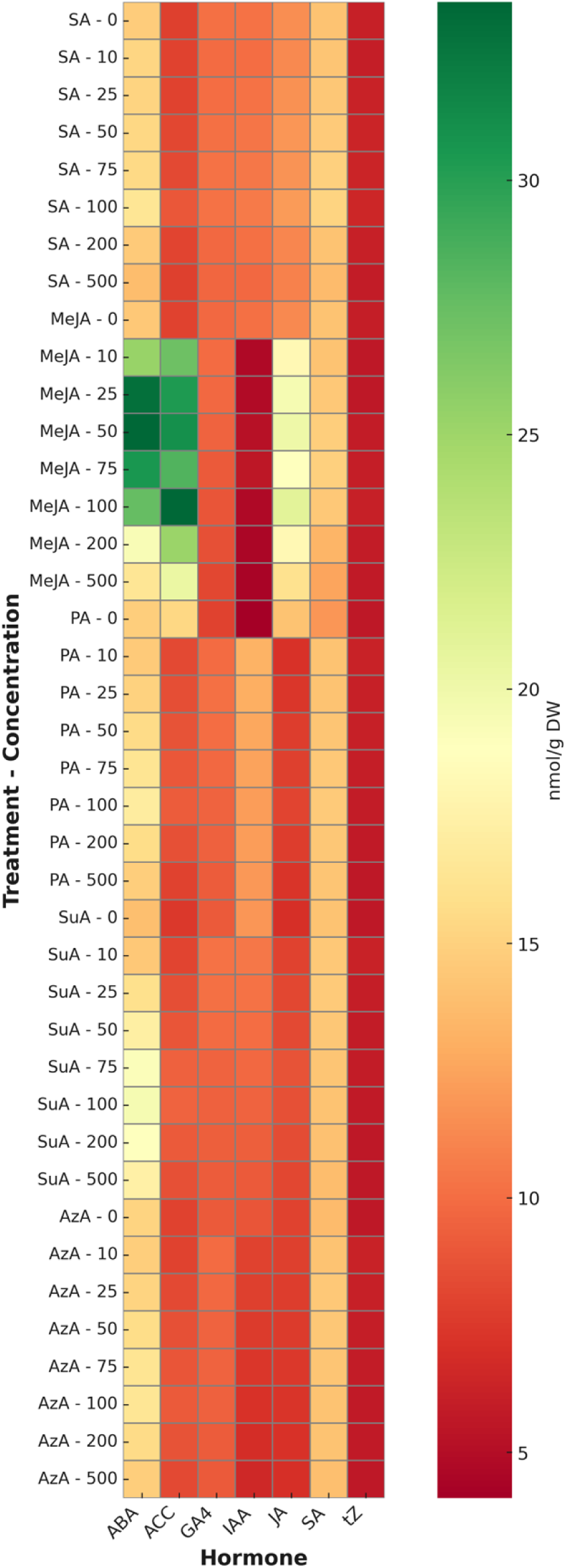
Heatmap of phytohormone concentration in *C. quitensis* in response to different elicitors. Concentration values were presented as nmol per gram of dry mass. Treatments correspond to SA, MeJA, PA, SuA, and AzA, applied at concentrations of 10, 25, 50, 75, 100, 200, and 500 µM. Phytohormones analyzed include ABA, ACC (ethylene precursor) GA_4_, IAA, JA, SA and tZ.

In contrast, IAA decreased under MeJA, reaching 4.7 nmol × g^−1^ DW at 100 µM. GA and tZ were also reduced in this treatment, with values of 2.8 and 1.5 nmol × g^−1^ DW, respectively (Fig. 2).

SA levels increased in SA treatments, reaching 6.4 nmol × g^−1^ DW at 50 µM, whereas ABA remained relatively stable, ranging between 13.0 and 15.4 nmol × g^−1^ DW from 10 to 50 µM. Within the same range, JA varied from 6.5 to 7.7 nmol × g^−1^ DW, and ACC ranged from 10.8 to 12.9 nmol × g^−1^ DW (Fig. 2).

PA induced intermediate ABA levels, peaking at 15.7 nmol × g^−1^ DW at 75 µM, and slightly elevated JA up to 8.1 nmol × g^−1^ DW. Conversely, IAA decreased from 13.2 to 12.2 nmol × g^−1^ DW between 10 and 100 µM, and GA declined from 7.6 to 6.9 nmol × g^−1^ DW over the same range.

In SuA treatments, ABA reached 19.6 nmol × g^−1^ DW at 100 µM, while JA remained between 7.5 and 8.7 nmol × g^−1^ DW. Growth-related hormones (IAA, GA, and tZ) remained relatively stable at low to intermediate levels, ranging from 6.2 to 9.8 nmol × g^−1^ DW (Fig. 2).

Finally, AzA exhibited more balanced hormonal profiles. ABA and SA peaked at 16.5 and 5.9 nmol × g^−1^ DW, respectively, at 75 µM. JA reached its maximum at 100 µM with 8.2 nmol × g^−1^ DW. IAA, GA, and tZ showed slight declines with increasing concentrations, reaching minima of 6.4, 3.8, and 1.8 nmol × g^−1^ DW, respectively, at 500 µM (Fig. 2).

### 3.3. Gene expression in response to elicitors

The expression of the analyzed genes exhibited variations depending on the type and concentration of the elicitor applied. In general, genes involved in the phenylpropanoid pathway, including *pal1*, *pal2*, *pal3*, *pal4*, *c4h*, *4cl*, *chs*, and *chi*, showed a significant increase in response to MeJA and PA at intermediate concentrations (50-100 µM), reaching expression levels between 8 and 10 times higher than the control. In contrast, SA and AzA induced more moderate responses, with increases at low concentrations followed by a reduction at higher levels. SuA displayed a more heterogeneous expression pattern, with variations depending on the specific gene and concentration applied (Fig. 3).

**Figure 3.**
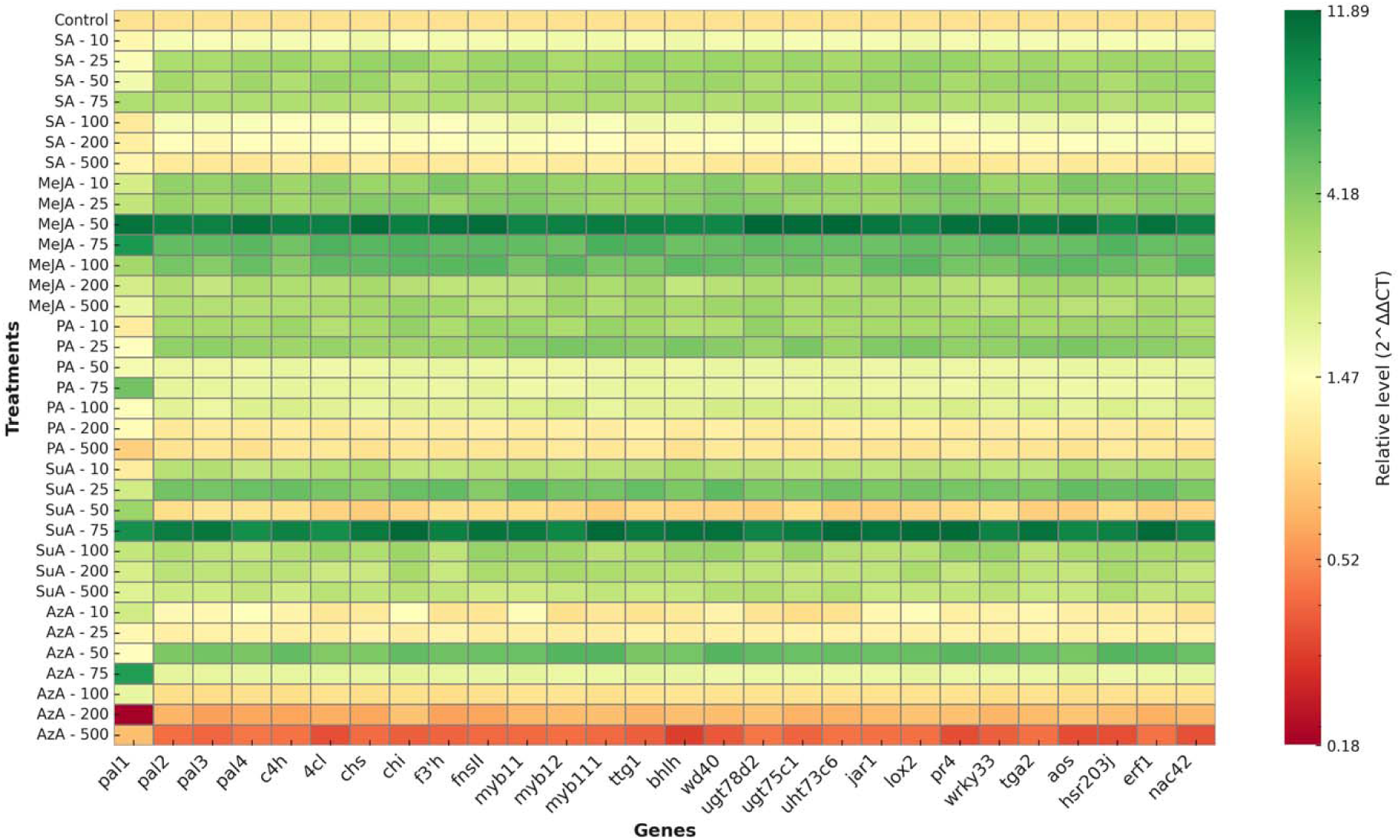
Heatmap of relative gene expression levels in *C. quitensis* in response to different elicitors. Expression values were presented as 2^−ΔΔCt^ relative to the control. Treatments correspond to SA, MeJA, PA, SuA, and AzA, applied at 10, 25, 50, 75, 100, 200, and 500 µM concentrations. Genes analyzed include those involved in the phenylpropanoid pathway (*pal1, pal2, pal3, pal4, c4h, 4cl, chs, chi*), flavonoid modification (*f3’h, fns ii, ugt78d2, ugt75c1, ugt73c6*), transcriptional regulation (*myb11, myb12, myb111, bhlh, wd40, ttg1*), stress metabolism (*wrky33, tga2, nac42*), jasmonate pathway (*lox2, aos, jar1*), and defense responses (*erf1, hsr203j, pr4*). Data represent the mean of three biological replicates.

Genes involved in flavonoid modification and conjugation, such as *fns II*, *ugt78d2*, *ugt75c1*, and *ugt73c6*, responded differentially to the treatments. MeJA and PA triggered the highest induction of FNS II, reaching expression levels over 8-fold higher than the control at 50-100 µM. Similarly, *ugt78d2* and *ugt75c1* exhibited peak expression under these elicitors, although with lower intensity, whereas *ugt73c6* showed a more uniform activation in the presence of SuA. In contrast, SA and AzA did not promote significant induction in these genes, and repression was even observed at higher concentrations.

Transcriptional regulators of the flavonoid pathway, *mybb11*, *myb12*, *myb111*, *bhlh*, *wd40*, and *ttg1*, displayed expression patterns like the structural genes of the pathway. Notably, MeJA and PA-induced sustained overexpression of *myb12* and *myb111*, while *mybb11* was more strongly activated in response to SuA. The *bhlh* and *wd40* factors exhibited increased expression under MeJA and PA, whereas *ttg1* responded primarily to SuA and PA in the 50-100 µM range.

Genes involved in stress metabolism regulation, such as *wrky33*, *tga2*, and *nac42*, exhibited elicitor-specific induction patterns. *wrky33* showed strong upregulation in response to MeJA and PA at 50-100 µM, whereas *tga2* displayed a more uniform increase under SuA treatment. *nac42* was predominantly activated by PA and SuA, reaching expression levels between 6- and 8-fold higher than the control at intermediate concentrations.

Genes associated with the jasmonate pathway, including *lox2*, *aos*, and *jar1*, responded primarily to MeJA, reaching expression levels up to 10 times higher than the control at 50-100 µM. SuA and PA also promoted their expression, although to a lesser extent, while SA and AzA did not induce significant changes in these genes.

Stress response and defense-related genes, *erf1*, *hsr203j*, and *pr4*, showed differential regulation depending on the treatment. *erf1* exhibited its highest expression in response to MeJA at 50 µM, whereas *hsr203j* responded more efficiently to SuA and PA at intermediate concentrations. *pr4* showed moderate activation with MeJA and SuA, while PA induced a more consistent response across all evaluated concentrations.

Overall, the results indicate that MeJA and PA at 50-100 µM are the most effective treatments for inducing the expression of key genes in the phenylpropanoid and flavonoid biosynthesis pathways, whereas SuA showed an intermediate effect, and SA and AzA elicited weaker and less consistent responses (Fig. 3).

### 3.4. Flavonoid production in response to elicitors

The accumulation of the flavones schaftoside, neoschaftoside, saponarin, and swertiajaponin in *C. quitensis* exhibited significant variations depending on the type and concentration of the elicitor applied. In general, intermediate concentrations of MeJA, PA, and SuA promoted the accumulation of these compounds, while higher concentrations (200-500 µM) tended to decrease their content compared to basal levels (Fig. 4).

**Figure 4.**
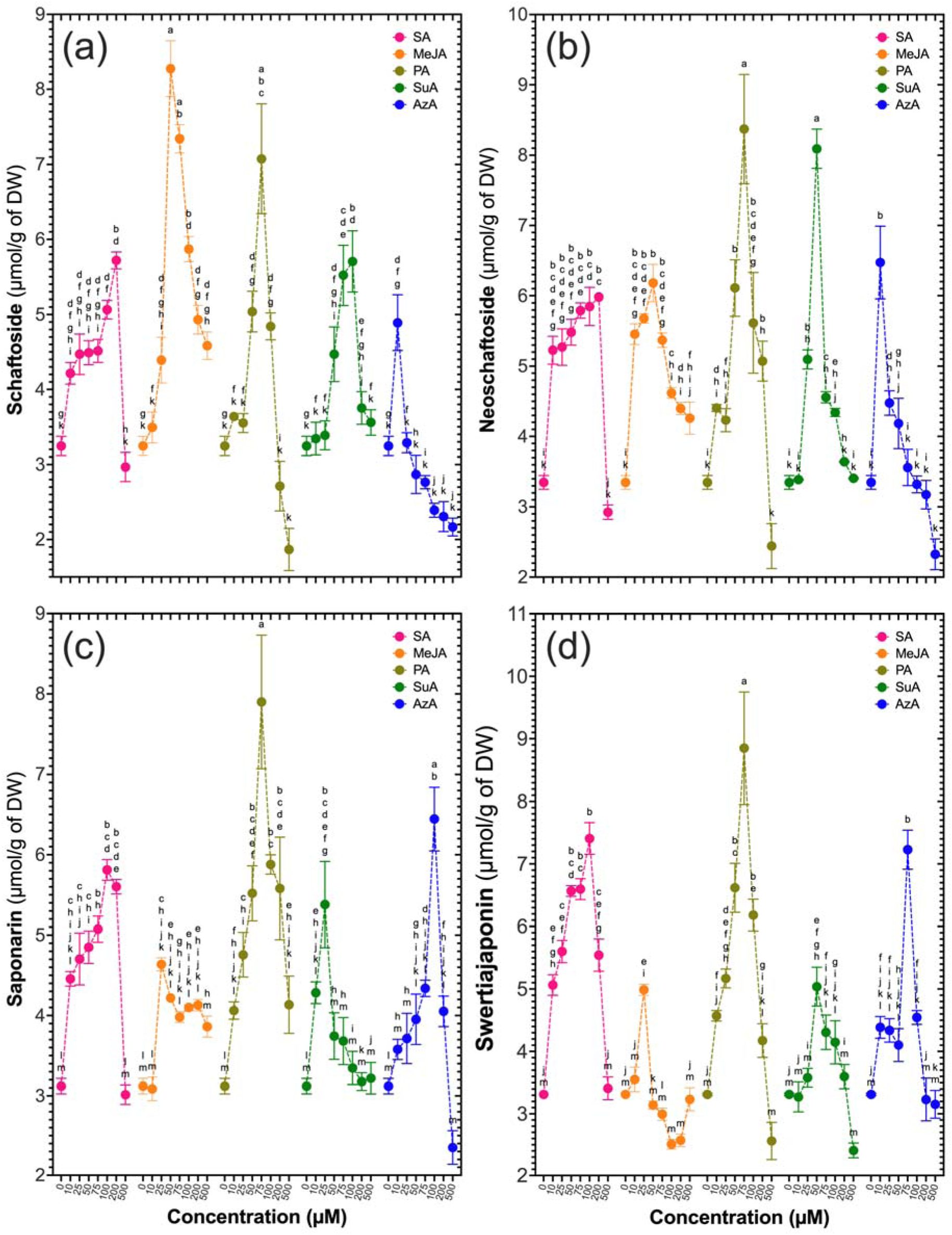
Effect of different elicitors on the accumulation of flavonoids in *C. quitensis*. (a) Schaftoside, (b) Neoschaftoside, (c) Saponarin, and (d) Swertiajaponin content expressed as µmol/g dry weight (DW) × yield. Treatments correspond to SA, MeJA, PA, SuA, and AzA at concentrations of 10, 25, 50, 75, 100, 200, and 500 µM. Data represent the mean ± standard error (SE) of three biological replicates. Different letters indicate statistically significant differences among treatments (*p* < 0.05, two-way ANOVA with Tukey’s post hoc test).

Schaftoside production displayed an induction profile dependent on the elicitor and concentration applied. MeJA and PA-induced the highest accumulation of this flavonoid, reaching values exceeding 8 µmol × g^−1^ DW at 50-100 µM. SuA also promoted a significant accumulation, with a sharper peak at 75 µM. In contrast, SA and AzA showed a more moderate induction, with a slight increase at low concentrations followed by a progressive reduction at higher levels (Fig. 4a).

Neoschaftoside accumulation followed a trend like that of schaftoside, with the highest values recorded in response to MeJA and PA in the 50-100 µM range, where concentrations of up to 10 µmol × g^−1^ dry weight were reached. SuA induced a moderate increase, while SA and AzA led to lower accumulation levels, with a decreasing trend at higher concentrations (Fig. 4b).

Saponarin production was strongly influenced by the elicitors, with a marked increase in response to PA and SuA, particularly at intermediate concentrations (50-100 µM). MeJA also promoted its accumulation, although with lower intensity compared to PA and SuA. Conversely, SA and AzA induced a more variable response, with increases at low concentrations but no clear pattern at higher concentrations (Fig. 4c).

Swertiajaponin accumulation exhibited a differential response depending on the elicitor. SA induced the highest increase in the production of this compound, reaching maximum values at concentrations of 50-100 µM. MeJA and PA also induced its accumulation, although to a lesser extent, whereas SuA and AzA resulted in lower levels compared to the other treatments (Fig. 4d).

Overall, these results indicate that intermediate concentrations of MeJA, PA, and SuA (50-100 µM) are the most effective in promoting the accumulation of schaftoside, neoschaftoside, and saponarin. In contrast, SA was more efficient in inducing swertiajaponin (Fig. 4).

### 3.5. Inhibition of *Botrytis cinerea* growth

The inhibition of *B. cinerea* growth exhibited a response dependent on the type of elicitor and the concentration applied in *C. quitensis*, revealing distinct inhibition patterns. MeJA showed the highest antifungal effect on derived extracts, reaching inhibition levels above 99% relative to the control in the 50-75 µM range. The inhibition was particularly pronounced in extracts from plants treated with 50 µM, followed by a gradual decline at higher concentrations. SA also promoted significant growth inhibition in extracts, with maximum values close to 72% at 25-50 µM. At higher concentrations (100-500 µM), the inhibitory effect of the application of SA in *C. quitensis* progressively decreased, suggesting a possible attenuation of its activity at elevated doses.

PA and SuA treatments exhibited intermediate effects in the extracts, with 63-72% inhibition levels at 50-100 µM applied concentrations in *C. quitensis*. The maximum inhibition for these treatments was observed at 75 µM, followed by a decline at 200-500 µM. The response pattern for PA showed more sustained inhibition across the tested concentration range. In contrast, SuA showed a more gradual increase in the inhibitory activity of extracts up to its peak at 75 µM, followed by a sharper reduction.

AzA was the elicitor with the lowest impact in the extracts derived from *C. quitensis* against *B. cinerea*, with significantly lower growth inhibition values compared to the other treatments. Although moderate increases in inhibition were observed at 50-75 µM, its effect was less pronounced and more variable. At concentrations above 100 µM, inhibition by AzA-treated plantlets declined sharply, suggesting that this compound does not exert a strong induction of antifungal compounds in *C. quitensis*.

Overall, MeJA at 50-75 µM and SA at 25-50 µM were the most effective priming treatments in *C. quitensis* to induce inhibiting compounds against *B. cinerea* growth. At the same time, PA and SuA exhibited moderate effects, and AzA showed the lowest induction of antifungal activity (Fig. 5).

**Figure 5.**
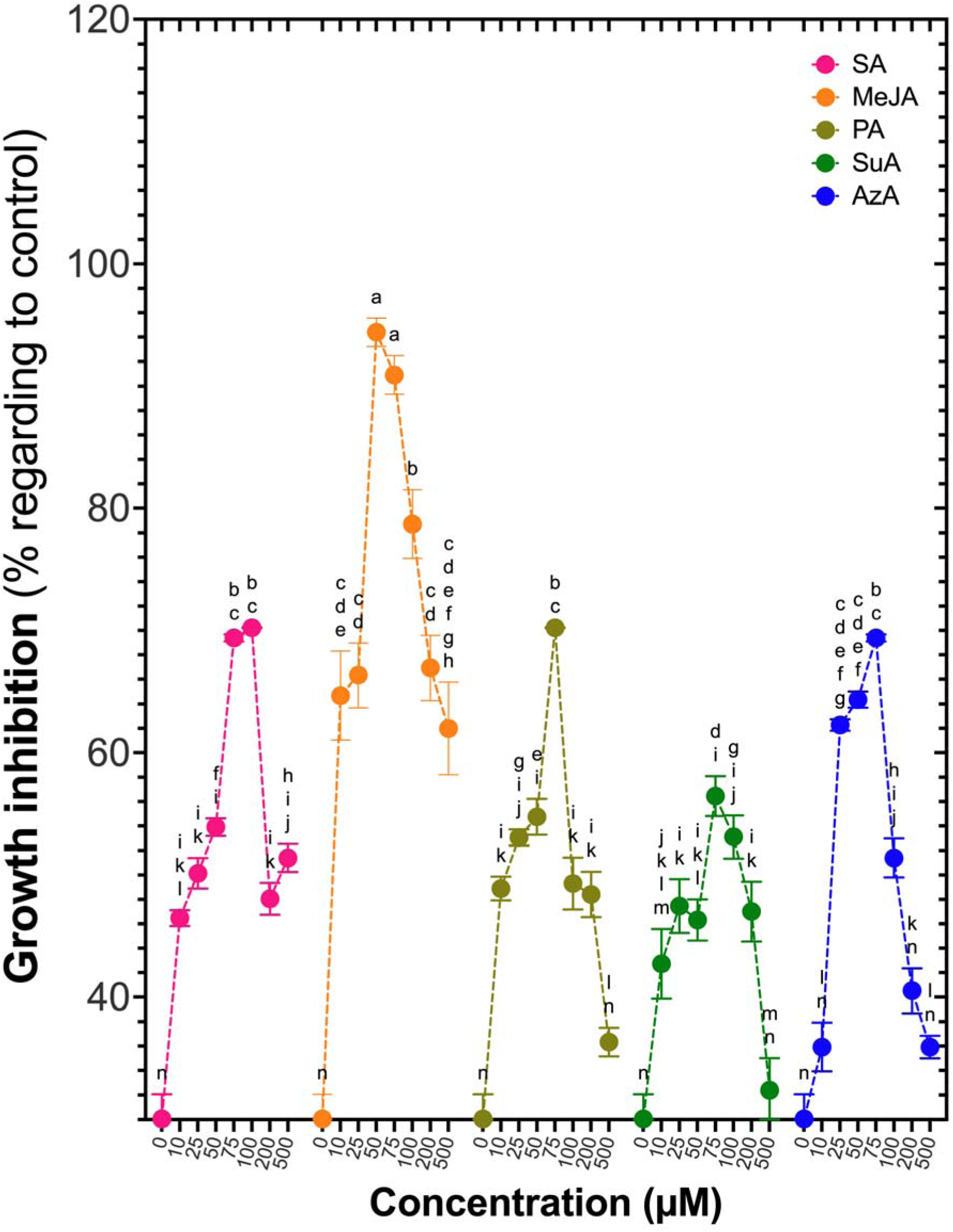
Effect of different elicitors on the inhibition of *Botrytis cinerea* growth. Growth inhibition is expressed as a percentage relative to the untreated control. Treatments correspond to SA, MeJA, PA, SuA, and AzA at concentrations of 10, 25, 50, 75, 100, 200, and 500 µM. Data represent the mean ± standard error (SE) of three biological replicates. Different letters indicate statistically significant differences among treatments (*p* < 0.05, two-way ANOVA with Tukey’s post hoc test).

## 4. DISCUSSION

The results of this study confirm that the exogenous application of the elicitors SA, MeJA, PA, SuA, and AzA modulates the accumulation of C-glycosyl flavones in *C. quitensis*, impacting both the phenylpropanoid metabolism and antifungal activity against *B. cinerea*. MeJA and PA were the most effective treatments for biomass induction, PAL activation, and flavonoid accumulation. In contrast, SuA had an intermediate effect, and SA and AzA exhibited more variable and less consistent responses. These findings align with previous studies demonstrating the role of MeJA and SA in activating plant defense pathways and promoting the synthesis of secondary metabolites with antifungal properties ^11,15,16^.

PAL activation is a key indicator of phenylpropanoid pathway modulation and is closely linked to flavonoid biosynthesis. In this study, PAL activity was regulated in a manner dependent on the elicitor type and concentration, with MeJA promoting the highest induction at 50 µM. This result is consistent with reports in *Arabidopsis thaliana* and other species, where MeJA has been shown to induce PAL activity and flavonoid accumulation through the regulation of transcription factors such as *myb12* and *myb111* ^17,18^. PAL activation was also significant under PA and SuA treatments, suggesting that these dicarboxylic acids may be involved in modulating phenylpropanoid biosynthesis, potentially through mechanisms related to oxidative stress and phenolic metabolite accumulation ^19,20^.

Hormonal profiling supported these observations, revealing strong increases in JA and ABA under MeJA treatments, with peak values of 20.9 and 27.6 nmol × g^−1^ DW, respectively, at 100 µM. ACC also increased markedly in the same treatment, reaching 33.4 nmol × g^−1^ DW, indicating activation of both jasmonate and ethylene-related defense pathways ^21^.

Gene expression analysis revealed that key genes in the phenylpropanoid and flavonoid biosynthesis pathways were differentially induced by the treatments. MeJA and PA promoted the upregulation of *pal* isoforms, *c4h*, and *4cl*, indicating an enhancement of the metabolic route leading to C-glycosyl flavone synthesis. These results align with findings in *Glycine max* and *Vitis vinifera*, where MeJA has been reported as a strong inducer of these pathways via the activation of *myb* transcription factors ^22–24^. Additionally, SuA stimulated the expression of *myb11* and *ttg1*, suggesting that this dicarboxylic acid may modulate flavonoid biosynthesis through transcriptional regulatory complexes. In contrast, SA and AzA induced a more heterogeneous and less pronounced response, consistent with previous reports where AzA functions primarily as a signaling molecule rather than a direct inducer of secondary metabolite accumulation ^25^ (Fig. 6).

**Figure 6.**
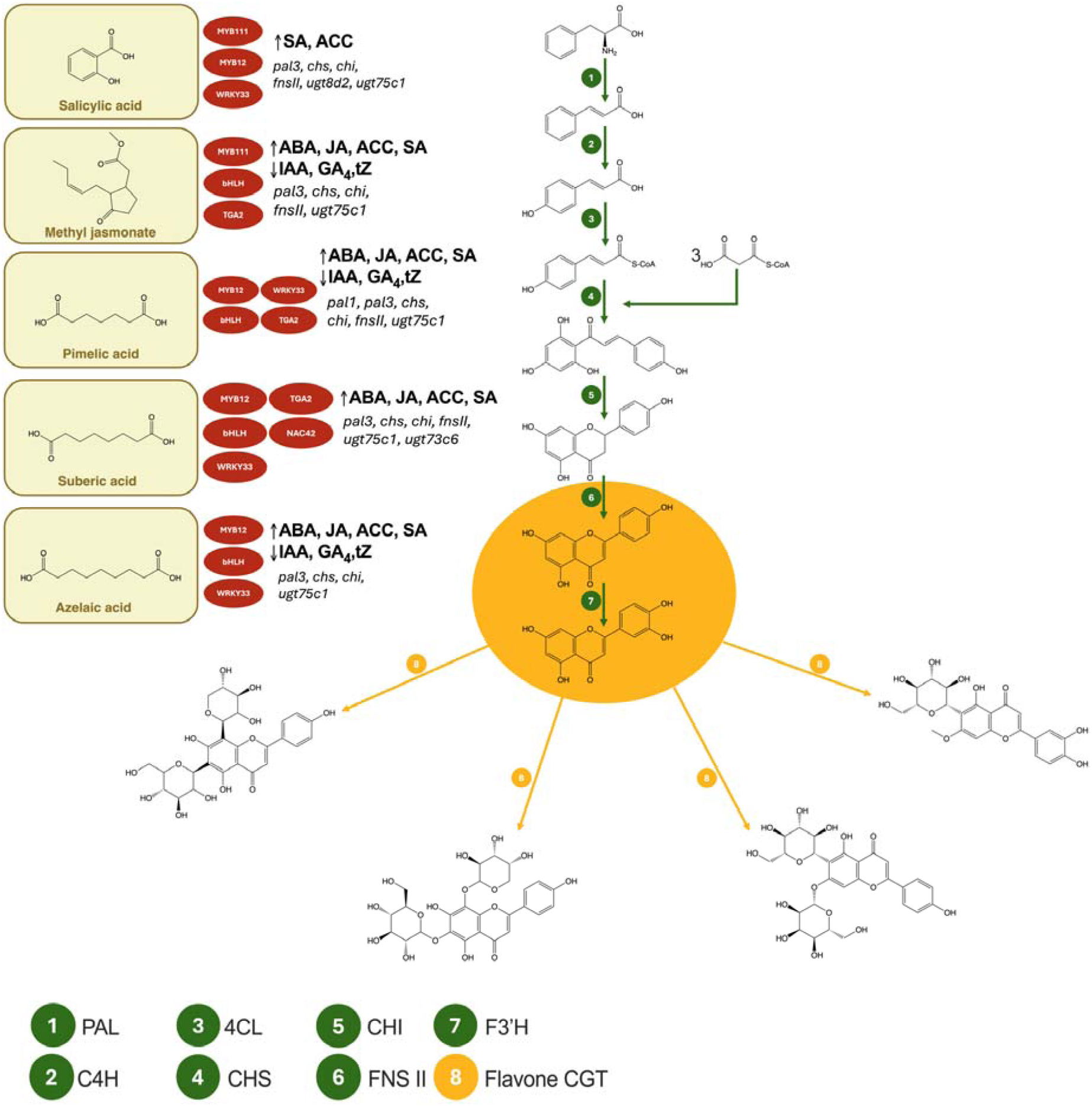
Schematic representation of the transcriptional regulation of flavonoid biosynthesis in *Colobanthus quitensis* in response to different elicitors. The left panel shows the chemical structure of the elicitors SA, MeJA, PA, SuA, and AzA, along with the transcription factors upregulated by each treatment. These include members of the MYB, bHLH, WRKY, TGA, and NAC families, which are known to regulate genes involved in the phenylpropanoid and flavonoid biosynthetic pathways. The right panel illustrates the flavonoid biosynthesis pathway, including PAL, C4H, 4CL, CHS, CHI, and FNS II enzymes, leading to the accumulation of C-glycosyl flavones. The genes induced by each elicitor are shown next to the respective transcription factors. This integrative model highlights how different signaling molecules differentially activate the transcriptional machinery involved in the accumulation of flavonoids with antifungal activity.

Concomitant with these gene expression trends, IAA, GA, and tZ levels decreased under SA and MeJA treatments, reaching 4.7, 2.8, and 1.5 nmol × g^−1^ DW, respectively, at 100 µM of MeJA. These declines in growth-related hormones are consistent with the observed reductions in biomass and support the metabolic shift towards secondary metabolism. In contrast, PA and SuA treatments maintained moderate levels of IAA (9.4–13.2 nmol × g^−1^ DW) and GA (5.4–7.6 nmol × g^−1^ DW), particularly at 10–75 µM, correlating with higher biomass accumulation and moderate PAL induction ^26^.

Regarding flavonoid accumulation, MeJA and PA were the most effective treatments for promoting the production of schaftoside, neoschaftoside, and saponarin at concentrations of 50-100 µM. At the same time, SA was more efficient in inducing swertiajaponin. These findings are consistent with studies in *Hypericum perforatum* and *Oryza sativa*, where MeJA and SA have been identified as key inducers of flavonoid biosynthesis ^27,28^.

The increase in secondary metabolites under MeJA and PA was paralleled by high endogenous JA levels (up to 20.9 nmol × g^−1^ DW in MeJA and 8.1 nmol × g^−1^ DW in PA), reinforcing the role of jasmonate signaling defense pathways in flavonoid induction. This resulted in the accumulation of secondary metabolites with antifungal activity ^29–31^.

The analysis of antifungal activity demonstrated that MeJA and SA were the most effective elicitors to induce antifungal compounds in *C quitensis* to inhibit *B. cinerea* growth, with maximum inhibition values of 99% and 72% at 50-75 µM and 25-50 µM, respectively. This is consistent with previous findings showing that MeJA and SA can induce plant defense responses, enhancing flavonoid and phenolic compound production with antifungal activity ^30^. Furthermore, the inhibition of *B. cinerea* growth suggests that the C-glycosyl flavonoids synthesized in response to elicitors may interfere with key fungal enzymes such as laccases, cellulases, and pectinases, a mechanism previously reported for similar compounds in other plant species ^10^.

Taken together, the hormonal data highlight a coordinated activation of defense-related pathways (JA, ABA, ACC) and repression of growth regulators (IAA, GA, tZ) under MeJA and SA treatments, explaining both the metabolic reprogramming and the antifungal outcomes observed ^32,33^.

The findings of this study have important implications for the scalability of bioactive metabolite production and its potential application in sustainable agriculture. The modulation of flavonoid biosynthesis through elicitors represents a viable strategy for developing natural biofungicides, which can be integrated into agricultural systems as an alternative to synthetic fungicides ^3,10,34^. The use of these compounds in organic farming is particularly relevant, as it enhances plant resistance to pathogens without compromising organic certification.

From a scalability perspective, the production of flavonoids in *C. quitensis* through elicitor treatments could be optimized in bioreactors or *in vitro* culture systems to maximize metabolite yields ^35^. Further studies should assess the economic feasibility of these processes, considering factors such as compound stability, production costs, and effectiveness under field conditions. Additionally, it would be relevant to evaluate the compatibility of these extracts with other integrated pest and disease management strategies for their implementation in diversified and sustainable agricultural systems.

## 5. CONCLUSIONS

This study demonstrates that the application of MeJA, PA, and SuA is an effective strategy for inducing flavonoid accumulation and enhancing antifungal activity in *C. quitensis* extracts against *B. cinerea*. The potential application of these findings in natural fungicide production and organic agriculture highlights the importance of further exploring the use of elicitors as key tools for crop protection and the development of sustainable agri-biotechnological products. Furthermore, the relatively low cost and broad availability of PA and SuA support their use in scalable biotechnological processes. When combined with controlled cultivation systems such as temporary immersion bioreactors, these elicitation strategies offer an efficient and sustainable pathway for the industrial production of high-value bioactive metabolites.

## AUTHOR CONTRIBUTIONS

**Lydia Rubilar:** Investigation, Methodology, Formal analysis, Data curation, Visualization, Writing – original draft.

**Javiera Avilés:** Investigation, Methodology, Formal analysis, Data curation, Visualization, Writing – original draft.

**Michelle Sarmiento:** Investigation, Methodology.

**Felipe Sobarzo:** Investigation, Methodology.

**Gustavo E. Zúñiga:** Supervision, Project administration, Funding acquisition, Resources.

**Rodrigo A. Contreras:** Conceptualization, Supervision, Project administration, Investigation, Formal analysis, Methodology, Validation, Writing – review and editing, Visualization, Resources, Funding acquisition.

## DATA AVAILABILITY

Data will be made available on request.

## ACKNOWLEDGEMENTS

This work was supported by the FONDECYT grant No. 3160274. The authors also wish to thank Dr. Richard Jeannotte and Dr. Pablo Zamora for their valuable collaboration in the initial metabolite identification analyses carried out at the University of California, Davis.

